# On the effect of asymmetrical trait inheritance on models of trait evolution

**DOI:** 10.1101/768820

**Authors:** Pablo Duchen, Michael L. Alfaro, Jonathan Rolland, Nicolas Salamin, Daniele Silvestro

## Abstract

Current phylogenetic comparative methods modeling quantitative trait evolution generally assume that, during speciation, phenotypes are inherited identically between the two daughter species. This, however, neglects the fact that species consist of a set of individuals, each bearing its own trait value. Indeed, because descendent populations after speciation are samples of a parent population, we can expect their mean phenotypes to randomly differ from one another potentially generating a “jump” of mean phenotypes due to asymmetrical trait inheritance at cladogenesis. Here, we aim to clarify the effect of asymmetrical trait inheritance at speciation on macroevolutionary analyses, focusing on model testing and parameter estimation using some of the most common models of quantitative trait evolution. We developed an individual-based simulation framework in which the evolution of species phenotypes is determined by trait changes at the individual level accumulating across generations and cladogenesis occurs then by separation of subsets of the individuals into new lineages. Through simulations, we assess the magnitude of phenotypic jumps at cladogenesis under different modes of trait inheritance at speciation. We show that even small jumps can strongly alter both the results of model selection and parameter estimations, potentially affecting the biological interpretation of the estimated mode of evolution of a trait. Our results call for caution when interpreting analyses of trait evolution, while highlighting the importance of testing a wide range of alternative models. In the light of our findings, we propose that future methodological advances in comparative methods should more explicitly model the intra-specific variability around species mean phenotypes and how it is inherited at speciation.

## Introduction

Understanding the mechanisms underlying the evolution of phenotypic traits is fundamental when addressing questions about functional diversity, ecological interactions, and the overall build-up of biodiversity (Futuyma and Agrawal, 2009; Katzourakis et al., 2009; Campbell and Kessler, 2013). Inferences about the processes governing phenotypic changes are facilitated by the use of macroevolutionary models, which deal with the evolution of phenotypic traits among related species (Stanley, 1979; Lande, 1980; Felsenstein, 1985, 2004). These macroevolutionary models are able to infer the evolution of a phenotype under a variety of scenarios including neutral or adaptive evolution (Lande, 1976; Beaulieu et al., 2012), as well as early and rapid phenotypic differentiation (Harmon et al., 2010; O’Meara, 2012). These models have been used to address a variety of evolutionary processes, helping us understand, for instance, body size evolution in mammals (Venditti et al., 2011; Slater, 2013), adaptation in *Anolis* lizards (Losos, 2011), evolutionary processes in extinct and extant vertebrates (Wagner, 2017; Silvestro et al., 2018), or changes in flower morphology and pollination syndromes (Wasserthal, 1997; Futuyma and Agrawal, 2009; Serrano-Serrano et al., 2017).

The simplest macroevolutionary model of trait evolution assumes that a trait *X* evolves following a random walk process, which is typically modeled by Brownian Motion (BM; Felsenstein, 1973). Under BM, the expected change in trait *X* is normally distributed and depends on time *t* and a rate parameter *σ*^2^, such that 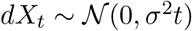, where the variance *σ*^2^*t* increases monotonically with time. We can also assume that traits evolve toward an optimal value and that phenotypic variance is constrained around that optimum, which leads to a process that is typically modeled by the Ornstein-Uhlenbeck (OU) model (Hansen, 1997). Many extensions of the BM and OU models have been developed and these include rate changes across clades (Eastman et al., 2011) or through time (Harmon et al., 2010), evolutionary trends (Slater et al., 2012; Silvestro et al., 2018), and trait boundaries (Boucher and Démery, 2016). Other extensions of BM are designed to identify bursts or “jumps” in the rate of evolution using Lévy processes. These approaches couple BM evolution with Poisson jumps (Landis et al., 2012; Duchen et al., 2017) to account for the presence of large changes in the rate of evolution that could capture shifts to new adaptive zones related to the dispersal into new geographic areas (as described by Simpson, 1944), key innovations, or by rapid climatic changes (Simpson, 1944; Hansen and Martins, 1996). The BM and OU models, and their extensions, all describe the anagenetic change of a trait as a continuous-time Markov chain along each branch of the tree, while the descendent nodes following a branching event inherit the last phenotypic value of the parent species (Felsenstein, 1973). In other words, at speciation, most current models of trait evolution assume that the descendent species inherit identically the phenotype of their last common ancestor (Felsenstein, 1973; O’Meara et al., 2006) leading to a symmetrical trait inheritance between the daughter lineages.

Macroevolutionary models use species as the units of evolution and aim to model the evolution of the mean phenotype of each species. This emphasis on a single trait value, which makes the commonly used models of trait evolution highly tractable, masks the fact that species are composed by sets of individuals possessing non-identical trait values (Mendes et al., 2018).

Because daughter populations after speciation are samples of a parent population (Fig. 1A), we can expect their mean phenotypes to differ from one another and from the parent mean due to random demographic events and/or differential segregation of the trait values within the incipient species (Uyeda et al., 2011; Kostikova et al., 2016). If the two mean phenotypes of the descendants differ from the parent phenotype, the difference will generate “jumps” of the mean phenotype at cladogenesis and lead to asymmetrical trait inheritance. Strong asymmetrical trait inheritance is expected when the trait is itself involved or even driving the speciation process (Fig. 1D), such as floral size and pollination syndromes in flowering plants (Wasserthal, 1997; Serrano-Serrano et al., 2015), morphology in insect genitalia (Schilthuizen, 2003; Ayoub et al., 2005), or color patterns in cyclids (Galis and Metz, 1998) and *Heliconius* butterflies (Jiggins et al., 2008). Even if the trait is not directly involved in the speciation process, geographic gradients of phenotypes and allopatric or peripatric speciation events could generate cladogenetic jumps in the mean phenotypes (Fig. 1B,C). For example, current subspecies of brown bear display substantial differences in their body mass (e.g. Kingsley et al., 1983; McCarthy et al., 2009). If they were to become independent species, the mean body size of each of the new species would differ from each other, and more importantly, would not be equal to the mean body mass of all brown bears. Geographic isolation of populations leading to speciation is likely to generate descendent lineages whose phenotypic means at the time of speciation differ from the overall mean of the parent species. For example, in the plant genus *Impatiens* it is hypothesized that habitat fragmentation and Pleistocene *refugia* triggered diversification, generating most of its current ca. 1000 species (Janssens et al., 2009) and their tremendous morphological diversity (Yu et al., 2016). Asymmetrical trait inheritance at speciation between isolated populations is likely to have played a role in shaping the phenotypic diversity of that clade. Therefore, at least in some cases, the assumption of identical inheritance might provide a poor description of the speciation processes.

**Figure 1:**
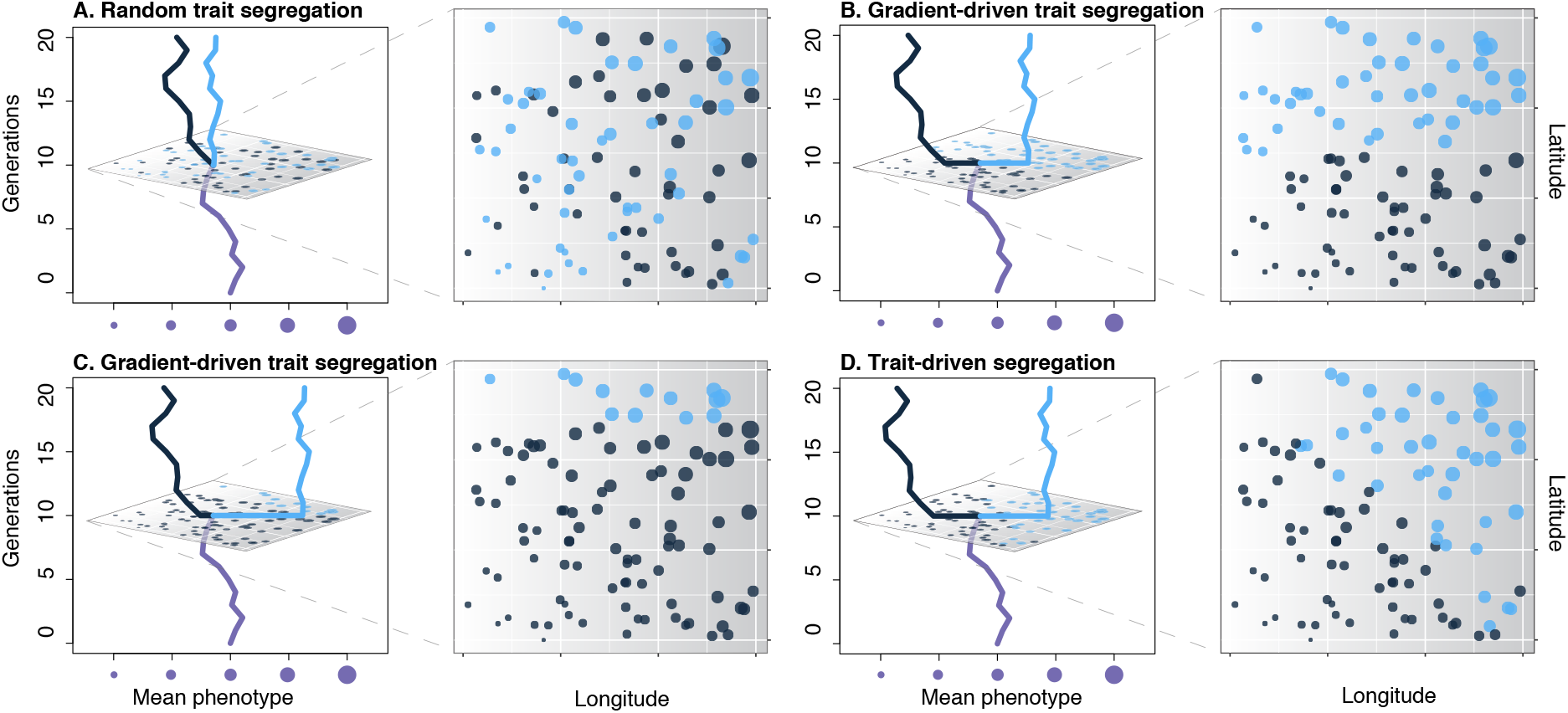
Four different scenarios of trait inheritance at speciation. Left panels: Evolutionary history of the mean phenotype before and after a speciation event. Right panels: Geographical distribution of the individuals at the speciation event depicted in the left panel. Bubbles represent the spatial distribution of individuals splitting from a parent population into two species (gray and light blue). The size of the bubbles is proportional to each individual’s phenotype. In this example, individual phenotypes follow a geographical gradient with smaller trait values on the lower-left part of the geographic range and larger trait values on the upper-right part. Under random segregation (A) speciation is not driven by the trait displayed here nor by geography and might result from a change in an uncorrelated trait, e.g. polyploidization. The mean phenotypes of the two descendants are expected to be equal with some stochasticity (resulting in slightly different means). Speciation occurring by geographic isolation, e.g. allopatric (B) or peripatric (C) may result in asymmetrical trait inheritance even though the trait itself is not the cause of speciation. Finally, if the trait is itself driving the speciation we can expect a full segregation of phenotypes at speciation (D). In the present study, we analysed cases A and D, which represent the extremes in terms of the size of cladogenetic jumps.

Models accounting for phenotypic jumps at speciation have been proposed. For instance, Pagel (1999) used a transformation of the elements of the phylogenetic variance-covariance matrix, called *κ*, to model cladogenetic jumps in the trait value by transforming the branch lengths of a phylogeny to modulate the importance of cladogenetic events versus anagenetic evolution. An alternative model introduced a Bayesian approach to estimate rates of both anagenetic and cladogenetic phenotypic evolution (Bokma, 2008). However, the empirical application of these models is limited and it remains unclear whether the models can capture the signal of asymmetrical trait inheritance at speciation.

Here, we aim to clarify the effect of different modes of asymmetrical trait inheritance at speciation on macroevolutionary analyses. We focus in particular on the fit of the models and the estimation of the specific parameters using four models of phenotypic evolution: BM, OU, Lévy, and *κ*. To accomplish this, we first developed an individual-based simulation framework in which the evolution of species phenotypes is determined by trait changes at the level of individuals accumulating across generations. Cladogenesis occurs by separation of two subsets of the individuals into new lineages. The inherited phenotypes of the descendent species is thus the mean of the individuals’ trait values and might differ from one another and from the parent phenotype. Using simulations, we assess the magnitude of expected phenotypic jumps at cladogenesis under different modes of trait inheritance at speciation. We show that these jumps can strongly alter both model selection and parameter estimations in all the widely used macroevolutionary models of trait evolution.

## Materials and Methods

To show the effect of asymmetrical trait inheritance on model fit and parameter estimation, we developed an individual-based forward simulator. This allows us to model the evolution of mean species phenotypes through the microevolutionary change across individuals within species and their segregation at speciation. We simulated trait values forward in time under two types of fitness landscapes: flat (where all individuals have the same fitness regardless of their trait value) and normal (where individuals with trait values closer to the optimum peak have higher fitness than other individuals, Fig. SI-B.1). We applied for each type of landscape different modes of speciation, which determine the partitioning of individuals from a parent species into two descendent species. The forward simulations generate changes at the generational time scale (the phenotypes of the individuals within each species) that translate into macroevolutionary processes when looking at the mean phenotype of species and their phylogenetic relationships resulting from the speciation process. Our simulations thus provide a framework linking micro with macroevolutionary processes for a quantitative trait.

### Modes of trait inheritance at speciation

We considered two modes of trait inheritance at speciation: “Random Segregation at Speciation” (RSS) and “Trait-driven Segregation at Speciation” (TSS). Under the RSS scenario, at each speciation event, the individuals of the parent population are randomly assigned to each daughter lineage (Fig. 1A). This corresponds to cases in which the phenotype represented by the trait of interest is completely independent of speciation. Conversely, under the TSS scenario, individuals of a parent population are split into two daughter lineages based on their phenotype and an arbitrary threshold defined a priori (Fig. 1D). The RSS and TSS constitute the extreme scenarios of how traits can be segregated during the split of a parent lineage and will generate the smallest and largest jumps, respectively. Cladogenetic jumps of intermediate size can also result from spatial gradients of phenotypic traits, captured by allopatric or peripatric speciation (Fig. 1B and C). To account for intermediate (more realistic) cases we also studied combinations or RSS and TSS, and we named them “mixed” scenarios. More precisely, we analyzed mixed scenarios where 5% and 50% of the cladogenetic events are TSS (and the rest are RSS).

### Death probabilities

Before the full description of the forward simulations (modes of reproduction, generational changes, etc.) and to unburden the explanation of the simulation algorithm, we will detail here the factors influencing the death probabilities of the simulated individuals.

The death probability of each individual (*δ*) is determined by the combined effect of three factors: global fitness, distance from the species-specific mean phenotype, and species-specific carrying capacity. Each of these factors contributes to the death probability of an individual by a specific quantity, such that

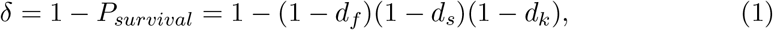

where *d_f_*, *d_s_*, and *d_k_* are the death probabilities by fitness, distance from the species-specific mean phenotype, and carrying capacity, respectively (see below).

#### Death probability determined by a global fitness landscape

We used a global fitness landscape to describe the fitness of individuals as a function of their phenotypic trait value (Simpson, 1944; Arnold et al., 2001; Svensson and Calsbeek, 2012; Boucher et al., 2017). The shape of the fitness landscape defines the survival probability of an individual and, as explained at the beginning of the Methods, we used two types of landscapes. First, we assumed that fitness was independent of the trait value, which corresponds to a flat landscape. Because the trait does not affect fitness under a flat landscape, its evolution is expected to follow a neutral process as captured by the BM model (Felsenstein, 1988). Second, we assumed a normally-shaped fitness landscape, where there is an optimal phenotype (the peak of the landscape, a global optimum that is independent of space) for which the fitness is highest. This landscape can represent the distribution of fitness under stabilizing selection (Beaulieu et al., 2012; O’Meara and Beaulieu, 2014), which is one of the biological processes resulting in an OU mode of evolution (Lande, 1976; Felsenstein, 1988; Hansen and Martins, 1996; Hansen, 1997).

In our simulations, the fitness landscape affects the death probability of each individual by an amount *d_f_*. Under a flat landscape, the death probability is independent of the trait value and set to a constant *d_f_* = *c* (Table 1). Under a normally-distributed landscape, the death probability of an individual depends on the position of the phenotypic trait value on the fitness landscape, which is described by an optimum *θ* and a variance 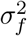. Thus, the fitness-based death probability of an individual with phenotype *x* is

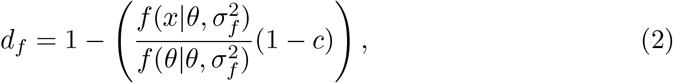

where 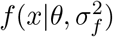 is the probability density function of a normal distribution with mean *θ*, and variance 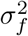 (a complete list of the parameters used in this study is given in Table 1). Under this formulation, the death probability of an individual with optimal phenotype (i.e. *x*_1_ = *θ*) is equal to *c*.

**Table 1:** Parameter definitions in our simulation framework. For each parameter we provide the fixed values used in simulations in which all species were assigned the same values, as well as the prior ranges from which parameters were drawn for each species in the variable-parameter simulations. Model parameters that are estimated from macroevolutionary models do not have any input value and are thus shown with a “not applicable” (*n.a.*) sign.

| Parameter | Definition | Fixed parameters | Variable parameters |
| --- | --- | --- | --- |
| $\mu$ | Root state | 0 | - |
| $\sigma_G^2$ | Parent-offspring variance | 0.01 | $\mathcal{U}(0.005, 0.02)$ |
| $c$ | Per individual baseline death probability (Eqs. (3),(2)) | 0.00001 | - |
| $\sigma_0$ | Initial standard deviation | 0.01 | - |
| $g$ | Growth rate | 1.2 | $\mathcal{U}(1, 2)$ |
| $\lambda_s$ | Speciation rate per lineage | 0.0001 | - |
| $K$ | Species specific carrying capacity | 2000 | $\mathcal{U}(1000, 4000)$ |
| $\sigma_f^2$ | Variance of the normal fitness landscape | 25 | $\mathcal{U}(10, 100)$ |
| $\sigma_s^2$ | Intraspecific variance | 1 | $\mathcal{U}(0.5, 5)$ |
| $\theta$ | Trait optimum | 5 | - |
| $\theta_{OU}$ | Trait optimum under OU | *n.a.* | *n.a.* |
| $\kappa$ | Statistic of the $\kappa$ model | *n.a.* | *n.a.* |
| $\alpha$ | Rate of mean reversion (or selective strength) | *n.a.* | *n.a.* |
| $\sigma_{BM}^2$ | Rate of evolution under BM | *n.a.* | *n.a.* |
| $\sigma_{OU}^2$ | Rate of evolution under OU | *n.a.* | *n.a.* |
| $\sigma_{LE}^2$ | Rate of evolution under Lévy | *n.a.* | *n.a.* |
| $\lambda$ | Jump rate | *n.a.* | *n.a.* |

#### Death probability determined by the distance to the species mean

We added a constraint to limit the intra-specific variance under the assumption that, regardless of the fitness landscape, there is selection against outliers diverging from the species’ mean (Harvey et al., 1991; Garland et al., 1993; Futuyma, 2010). To achieve this, we set the death probability *d_s_* of an individual with phenotype *x* to a function of its species mean phenotype at time *t* (*m*(*t*)) and of a fixed intra-specific variance 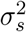. Thus, this death probability is

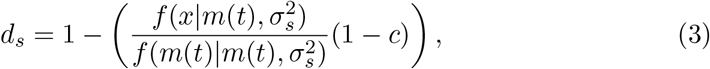

where 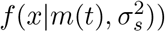 is the probability density function of a normal distribution with mean *m*(*t*), and variance 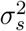 (Table 1). While the intra-specific variance is assumed to be constant over time, the species mean, *m*(*t*), is not subject to any constraints but simply computed as the empirical mean of the phenotypes of the individuals.

#### Death determined by the carrying capacity

Finally, the population growth within a species is controlled by a species-specific carrying capacity *K* (Etienne et al., 2011). This is achieved by changing the death probability *d_k_* of all individuals within a species based on their current population size *n* and its proximity to *K*, such that

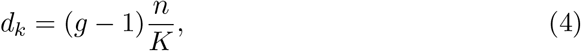

where *g* is the growth rate of that species.

### Forward simulations

For all combinations of fitness landscapes and modes of trait inheritance (RSS flat, RSS normal, mixed 5% TSS flat, mixed 5% TSS normal, mixed 50% TSS flat, mixed 50% TSS normal, TSS flat, and TSS normal), we ran an individual-based forward simulator of continuous traits which works as follows:

1. **Initialization.** At generation *i* = 1, we have a starting population of *n* = 200 individuals assigned to Species 1, with trait values drawn from a normal distribution with mean *µ* and standard deviation *σ*_0_ (Table 1).
2. **Change from one generation to another.** Individuals of generation *i* = 1 are replaced by a number of offspring based on a Poisson distribution *n*_*i*+1_ ~ *Pois*(*g* × *n_i_*), where 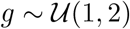 (Table 1). The trait values assigned to each individual of the new generation will be normally distributed with mean set to the value of the parent individual and variance 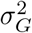. Individuals of the new generation die with probability *δ* (Eq. (1)). Based on this birth-death equilibrium, the overall population might go extinct or grow until a carrying capacity *K* is reached (see section *Death determined by the carrying capacity*). The variance of the normal landscape 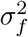 was set to a default value of 25 (and varied between 10 and 100) to ensure that the initial population will not start at a too-low fitness position (Table 1). This landscape, with its default value, yields a 1.25% increase in death rate for a phenotype = 0 (the predefined root state) compared to the death rate at phenotype = 5 (the predefined trait optimum).
3. **Speciation.** Speciation occurs with probability *λ_s_* per generation per lineage. At speciation, a given number of individuals from the parent species are assigned a new species label. The initial number of individuals for the new species is drawn from 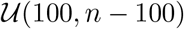, where *n* is the population size of the parent species. We note that speciation leads to a temporary reduction in population size in the two daughter lineages, but with the growth rate being always positive (Table 1), we ensure that the populations grow back rapidly until the carrying capacity is reached again within a few generations (Fig. SI-B.5). The way individuals are chosen to become a new species is described in the previous section *Modes of trait inheritance at speciation*.
4. We repeat steps 2 and 3 for every species for a total of 100,000 generations or until 100 species are generated. We then output the following files and statistics: species phylogenetic tree in newick format, mean (and variance) trait values per species at the tips of the phylogeny, mean (and variance) trait values per species at intermediate generations along the tree, full trait values along the entire phylogeny (optional setting), population sizes per species, species specific parameters, and mean (and variance) phenotypic-trajectory plots.

For each scenario (RSS flat, RSS normal, mixed 5% TSS flat, mixed 5% TSS normal, mixed 50% TSS flat, mixed 50% TSS normal, TSS flat, and TSS normal), we simulated 100 data sets, from which we extracted the species level phylogenies and trait values for further analyses. We summarized the sizes of jumps generated under the different simulation scenarios by computing the absolute difference between the phenotypes of the two daughter lineages following each speciation event.

### Fitting macroevolutionary models

To quantify the effect of the different simulation settings on species-level analyses of trait evolution, we analyzed the simulated phylogenetic trees (re-scaled to a root age of 1) and phenotypic values using the function *fitContinuous* from the R library *geiger* (Pennell et al., 2014) under different macroevolutionary models in a maximum likelihood framework. We fit the models BM, OU, *κ*, and we added also a white noise (WN) model, which indicates the absence of phylogenetic signal in the trait values. From all these models we calculated their respective AICc’s and parameter estimates (Table 1). Additionally, we analyzed the simulated data under a Lévy (LE) jump model using the software *levolution* (Duchen et al., 2017) and obtained the respective AICc and model parameters *σ_LE_* and jump rate *λ*. Since there is no implementation of the Bayesian/MCMC model by Bokma (2008), we did not include it in the current analysis. However, given that our simulations comprise mostly complete phylogenies and complete trait data, it is expected that the *κ* model will capture this signal equally well (Harmon, 2018).

### Parameter estimation

We then assessed the effect of the different speciation scenarios on the estimation accuracy of the following macroevolutionary parameters: the rate of evolution under BM 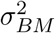, OU 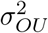, and Lévy 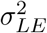, the jump rate *λ*, the *κ* parameter, the selective strength *α*, and the optimum trait value *θ_OU_*. For this purpose we re-run simulations but this time fixing parameters throughout the simulation, so that they don’t change along a phylogeny (Table 1, fixed parameters). To quantify the accuracy of the estimated 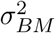 and 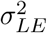 we ran the simulator again for 10,000 generations and calculated the true evolutionary rate 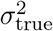 as the average change in the trait value per generation along a lineage. We then computed the relative error of the estimates as 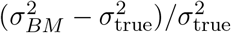. To quantify the accuracy of the estimated selective strength *α* we compared it to the true selective strength *α*_true_ using the estimator of Duchen et al. (2020), which takes as input the vector of mean phenotypes over time for each species. Finally, we computed the accuracy of the optimum trait value as (*θ_OU_* − *θ*)*/θ*.

## Results

We studied the effect of asymmetrical trait inheritance on the fit of macroevolutionary models of phenotypic evolution. Our individual-based simulations generated species trees reflecting mostly pure-birth processes, as shown by the lineage-through-time (LTT) plots generated for all scenarios (Fig. SI-B.2). The parameter values used in our simulations produced few extinction events, ranging from zero to three extinctions per simulations in RSS scenarios, and from zero to six in TSS scenarios (Table SI-A.1).

### Phenotypic trajectories

The simulated phenotypic trajectories show differences between different scenarios. The first clear distinction lies between flat and normal landscapes. Flat landscapes produce traits evolving without a trend and generate BM-like phenotypic trajectories (Fig. 2A, B). In contrast, with normal landscapes species mean phenotypes evolve toward the value with maximum fitness (Fig. 2C, D), generating an OU-like process (Revell, 2012).

**Figure 2:**
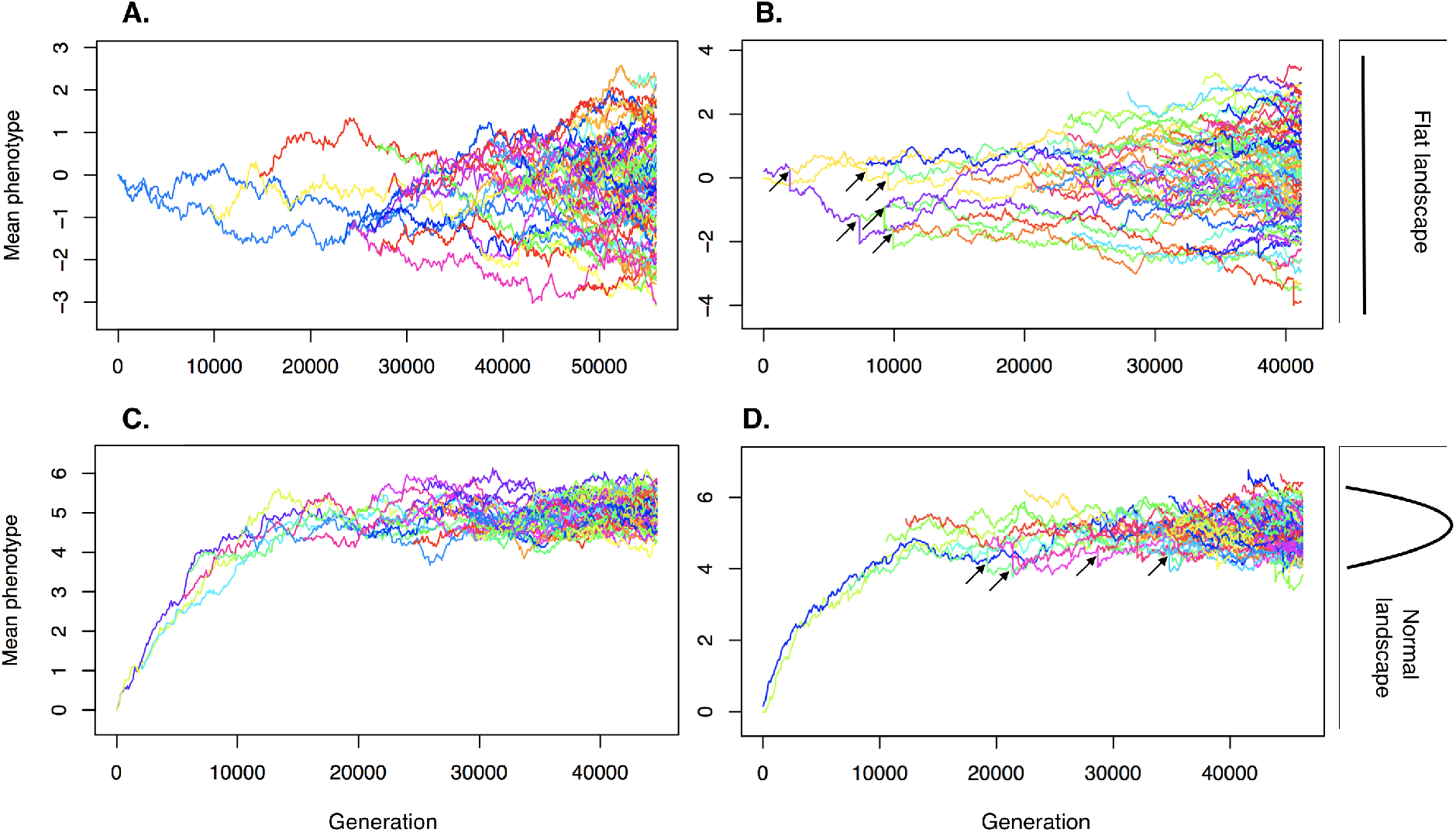
Examples of phenotypic trajectories under the extreme scenarios: A) RSS-flat, B) TSS-flat, RSS-normal, and D) TSS-normal. Different colors represent different species. Some examples of cladogenetic jumps (for scenarios B and D) are pointed with black arrows. The bulk of the jumps are covered by the lineage lines, but a quantification of their sizes is shown in Fig. 3.

When individuals were assigned to new species independently of the trait value (RSS scenarios), the trait inheritance at speciation is nearly symmetrical, as shown by the lack of noticeable jumps in the phenotypic trajectories (Fig. 2A, C). Conversely, if individuals were assigned to new species based on a threshold in the trait value (TSS scenarios), the trait inheritance at speciation is asymmetrical and results in visible jumps in species phenotypes (Fig. 2B, D). The jump sizes between the daughter species under our simulation settings were on average 24 times higher in the TSS scenarios compared to the RSS scenarios (Fig. 3), while mixed scenarios had intermediate jump sizes. Phenotypic jumps derive from the stochastic separation of individuals of the ancestral species into the two descendent lineages and will thus differ across speciation events in the phylogeny. The pattern of jump sizes was similar across flat and normal fitness landscapes (Fig. 3).

**Figure 3:**
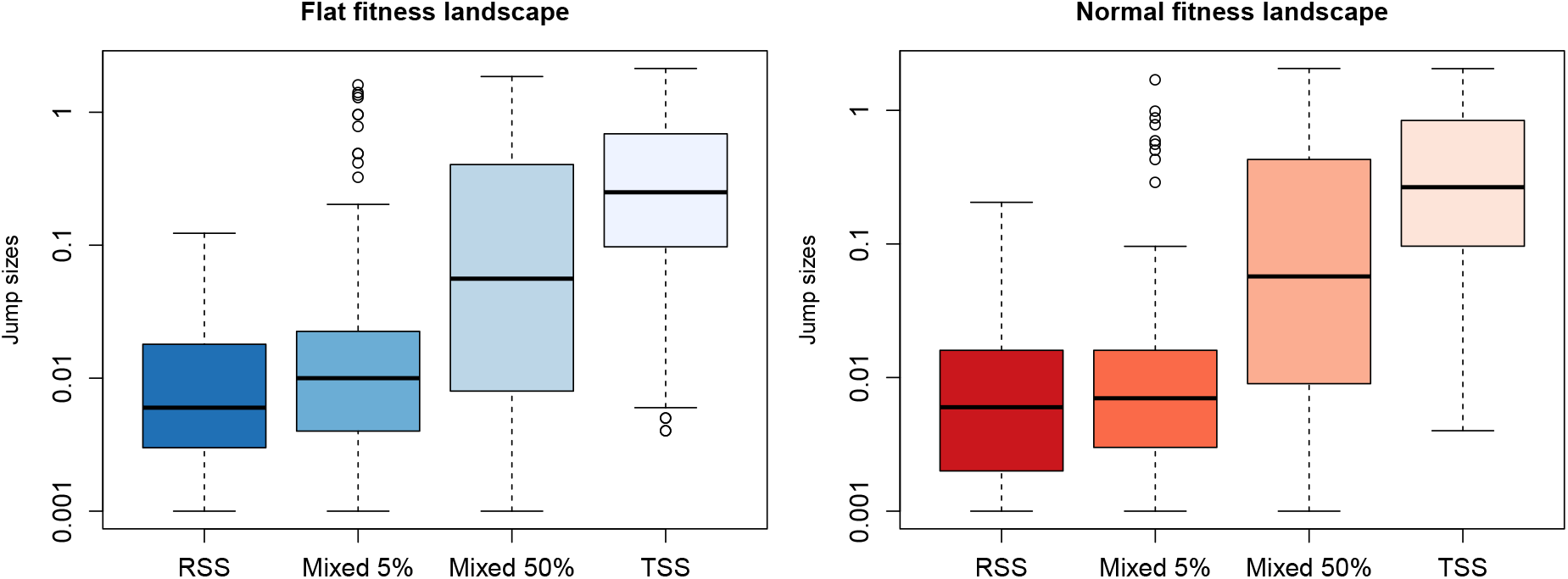
Absolute jump sizes for every scenario. Boxplots represent the distribution of jump sizes for all speciation events across all simulations for a given scenario. Here, the size of a jump is the absolute difference between the starting values of the two daughter lineages right after a cladogenetic event. Scenarios following a flat and normal fitness landscapes are colored following the color pattern shown in Fig. 4.

### Effect of RSS and TSS on model fitting

To analyze the effect of the different modes of trait segregation at speciation and fitness landscapes on model fit, we fitted BM, OU, Lévy, *κ* and WN models to the simulated trees and tip trait data. In general, for both flat and normal fitness landscapes, an increasing number of cladogenetic jumps (from RSS to TSS) affected model fit substantially (Fig. 4), an effect that cannot be attributed to consistent differences in the simulated trees (Table SI-A.2).

**Figure 4:**
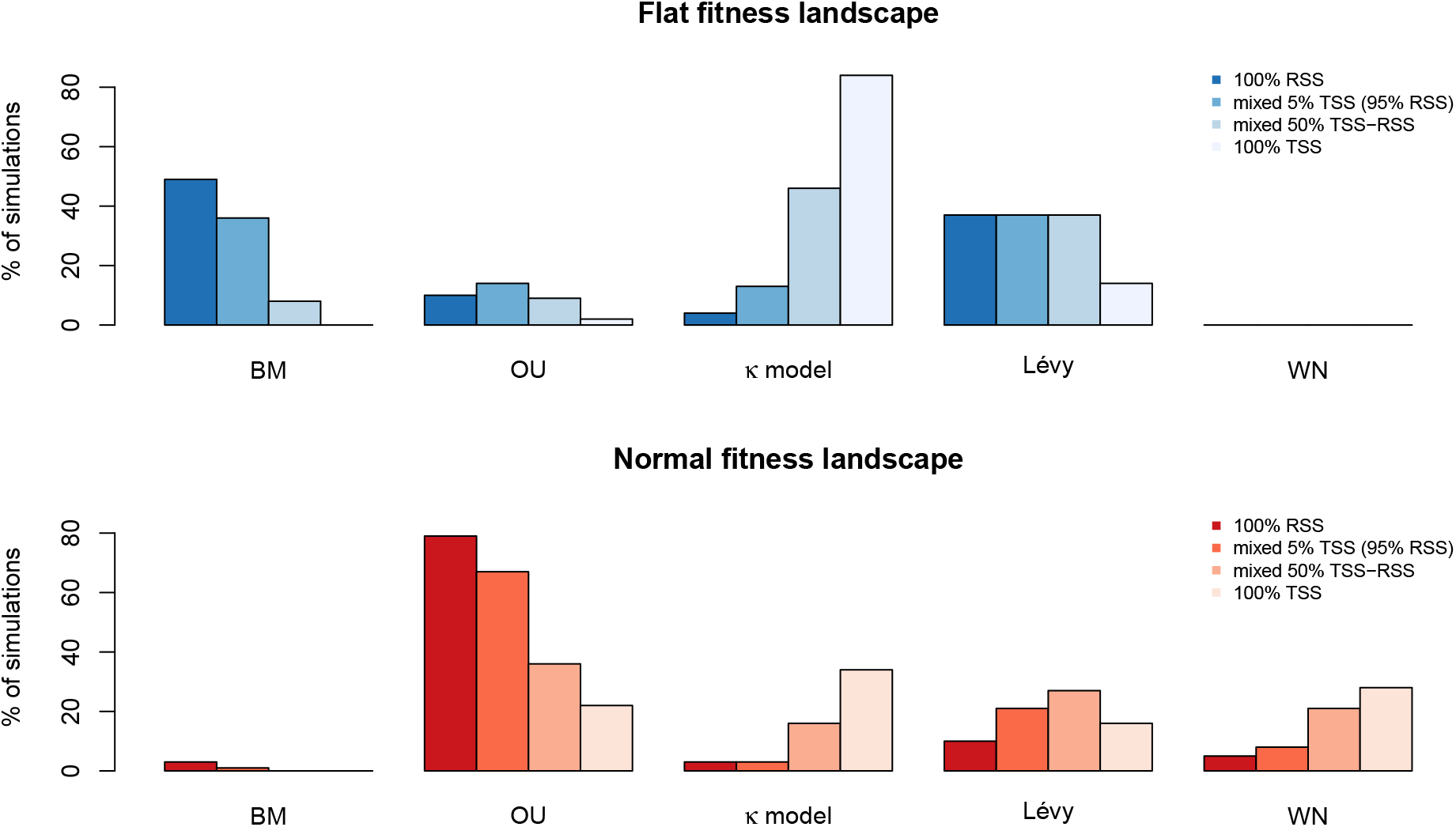
Model fit under all scenarios. Bars represent the percentage of simulations that were best fit by a given model. Parameters used for these simulations where drawn from uniform priors (Table 1, prior). Mixed scenarios consist of 5, and 50% of TSS cladogenetic events along a single simulation run. Abbreviations: BM = Brownian Motion, OU = Ornstein-Uhlenbeck, WN = White Noise. The second best fit is given in Fig. SI-B.4.

Under the flat fitness landscape the BM model was the most frequently selected for the RSS scenario, while the Lévy model was the best fit under mixed RSS/TSS scenarios, followed by BM or *κ* (Fig. 4, upper row). Concerning the full TSS the *κ* model was the most frequently selected, followed by Lévy (Fig. 4, upper row). Under normal fitness landscape, RSS and mixed 5%,10% scenarios had OU as the best fitting model, followed by Lévy. TSS and mixed 50% TSS scenarios had *κ* and Lévy as the most selected models, followed by WN (Fig. 4, lower row).

We performed one additional test where we set the death probability by its distance to the species mean *d_s_* to zero to check the influence of the *d_s_* parameter on model testing. We tested this on the RSS-flat scenario. We found that jump sizes under these circumstances (Fig. B.3, left) are on average larger than those under RSS where *d_s_* is taken into account (Fig. 3, upper row). On the other hand, we did not observe an increase in cases where BM is the preferred model (Fig. B.3, right).

### Effect of RSS and TSS on parameter estimation

Parameter estimation was performed on a new set of simulations with fixed parameters that do not change along a phylogeny (Table 1, fixed parameters). Analyses of data sets simulated under a flat RSS scenario resulted in a slight overestimation of the evolutionary rates under both the BM and Lévy models (Table 2). The overestimation was 50% lower under the Lévy model (Fig. 5A and 5B). This indicates that the Lévy model was able to identify, at least to some extent, the jumps generated under asymmetrical segregation at speciation. Simulations obtained from the flat TSS scenario resulted in a 2.58-fold overestimation of the evolutionary rate under BM and a 1.99-fold overestimation under Lévy (Table 2).

**Figure 5:**
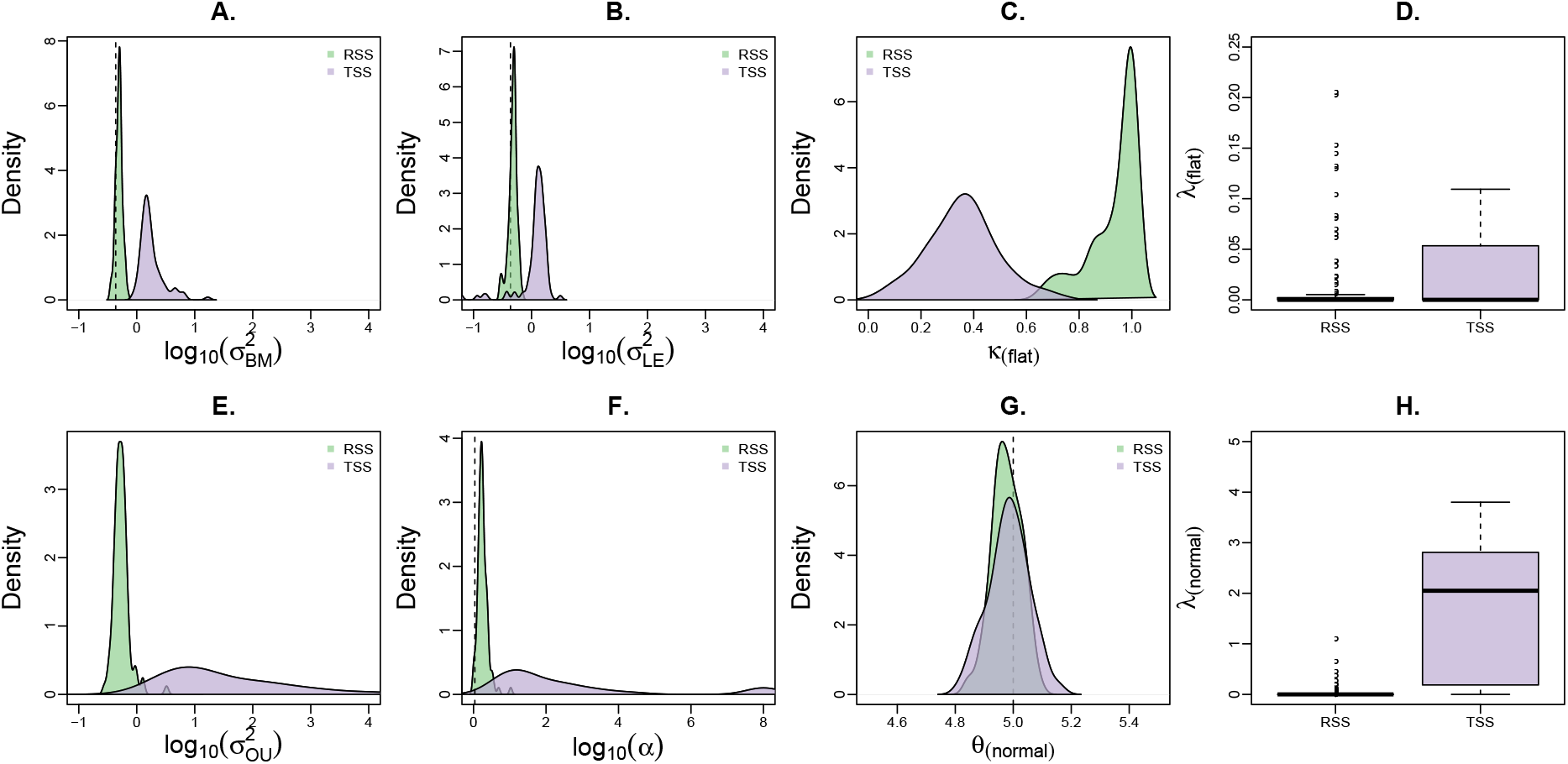
Parameter estimates of *fitContinuous* (R package *geiger*) of various model parameters. Green (purple) distributions represent the estimates of all TSS (RSS) simulations. Abbreviations: *σ*^2^=evolutionary rate, *κ*=parameter kappa, *α*=selective strength, *λ*=jump rate, and *θ*=optimum. Median relative error (MRE) between RSS or TSS and their true values are depicted in Table 2.

**Table 2:** Median relative error (MRE) of some parameter estimates.

| Parameter | MRE (RSS) | MRE (TSS) |
| --- | --- | --- |
| $\sigma_{BM}^2$ | 0.14 | 2.58 |
| $\sigma_{LE}^2$ | 0.12 | 1.99 |
| $\theta_{OU}$ | -0.004 | -0.002 |
| $\alpha$ | 0.57 | 29.79 |

In simulations with a normal landscape, the trait optimum was accurately recovered under all scenarios (Fig. 5G; Table 2), that is regardless of the segregation scenario at speciation. The strength of selection *α* of the OU process was only slightly overestimated under the RSS scenario, while it was strongly overestimated (29-fold) under the TSS scenario (Fig. 5F; Table 2).

Although for other parameters we could not compute accuracy (as the true values could not be derived analytically or empirically from the simulations), the estimated values differed vastly between speciation modes (Fig. 5), indicating that segregation at speciation has a large effect on the macroevolutionary modeling of trait evolution. For instance, the *κ* parameter was lower in TSS when compared to RSS (Fig. 5C), which is expected since *κ* values tend to zero when most of the variation is accumulated at speciation events (Pagel, 1999). Finally, the jump rate of the Lévy model (*λ*) was much higher in TSS than in RSS simulations, as expected given the increased amount of phenotypic jumps at speciation (Figs. 5D and 5H).

## Discussion

We explored the effects of asymmetrical trait inheritance on macroevolutionary analyses using phylogenetic comparative methods. We modeled the asymmetry of trait inheritance at speciation as a result of the segregation of individuals from a parent population into two daughter lineages. We showed that even small phenotypic jumps at cladogenesis can have profound effects on both model testing and parameter estimation using widely-used macroevolutionary models, most of which implicitly assume identical trait inheritance at speciation (Felsenstein, 1988; Hansen and Martins, 1996).

### A simulation framework

We built a simulation framework that captures some of the main evolutionary processes that affect the micro- and macroevolutionary history of a quantitative trait. By tracking the phenotype of each individual for thousands of generations and across all the branches of a phylogenetic tree, we studied the effect of asymmetrical trait inheritance at speciation on model fit and parameter estimation of traditional macroevolutionary models. The current version of our simulator is set to work with haploid taxa, two fitness landscapes: flat and normal, and three modes of trait inheritance at speciation: random segregation (RSS), trait-driven segregation (TSS), and mixed RSS/TSS. Extensions to diploid taxa, as well as various fitness landscapes and additional speciation modes will open a vast realm of potential applications. For instance, such a simulator can help answer longstanding questions about the impact and fate of microevolutionary processes on a macroevolutionary scale (Hansen and Martins, 1996; Arnold et al., 2001; Reznick and Ricklefs, 2009; Kostikova et al., 2016; Rolland et al., 2018; Duchen et al., 2020). Currently, our community is in need of individual-based simulators of phenotypic traits, although some research in this direction has been done in the past (Neuenschwander et al., 2008; Boucher et al., 2014; Aguilée et al., 2018; Polly, 2019). For instance, Neuenschwander et al. (2008) have developed a simulator of quantitative traits but they focus more on the genetic basis and works only at the population level. Additionally, Boucher et al. (2014) have also developed a simulator of phenotypes allowing for different modes of speciation, but limited to a small number of generations. Polly (2019), on the other hand, developed computational models that take spatial processes into account and applied them to paleontological data. His results also support the fact that overlooking such processes results in incorrect interpretations of traditional model fitting in Evolutionary Biology (Polly, 2019). Our simulator is unique in that it simulates populations of individuals for thousands of generations, and it allows for different types of speciation events to happen, under different adaptive peaks, thus generating complete phylogenies.

### Asymmetrical trait inheritance

Traditional macroevolutionary models (including BM and OU), when run along phylogenetic trees, assume that the last phenotypic value of a lineage and the starting values of its two daughter lineages are identical (Felsenstein, 1973). However, we showed that when we consider species as set of individuals each bearing its own phenotype, the resulting trait inheritance at speciation will no longer be identical, and there are plenty of examples in nature where this is the case (Galis and Metz, 1998; Lachaise et al., 1988; Schilthuizen, 2003; Coyne and Orr, 2004; Ayoub et al., 2005; Jiggins et al., 2008; Givnish et al., 2009; Serrano-Serrano et al., 2015). Consequently, when the segregation of individuals between daughter species is random with respect to the trait value (our RSS scenario), the expected amount of phenotypic change is small, although greater than zero (Fig. 3). Yet, our simulations showed that even these small deviations from the assumption of identical inheritance generate a substantial bias in the parameter estimates under standard evolutionary models and alter the robustness of model selection (Figs. 4, 5). While asymmetric inheritance has been theorized and modeled explicitly for discrete traits (Goldberg and Igić, 2012), and biogeographic ranges (Ree et al., 2005; Ree and Smith, 2008; Goldberg et al., 2011), this has not been formally explored for quantitative traits (but see Bokma (2008)).

### Effects of RSS

RSS generates jumps at lineage splits due to sampling effects (Fig. 3). We found that even very small deviations from the parent phenotype can alter the results of model fit (Fig. 4) and generate biased parameter estimations (Fig. 5). 50% of the datasets simulated under a flat landscape and RSS supported BM as the best model (Fig. 4, Flat landscape, BM). We found the same pattern even after removing the constraints on instraspecific variance *d_s_* = 0 (Fig. B.3). This scenario is supposed to fulfill all the requirements for BM (equal fitness, no boundaries, and almost symmetrical inheritance from parent to daughter species; Felsenstein, 1988; Boucher and Démery, 2016). However, we show here how the effect of random sampling of individuals generates small variation in the mean phenotypes, and is enough to violate model assumptions leading to the rejection of BM in half of the simulations. Conversely, the RSS-normal scenario is consistent with the expectations of an OU model (normal fitness landscape and almost symmetrical inheritance from parent to daughter species; Hansen, 1997, but see Cooper et al. 2016) and indeed OU was the model of choice in most simulated datasets under this scenario (Fig. 4, normal RSS).

### Effects of TSS

Bigger deviations from the parent mean (as expected for TSS but also for allopatric and peripatric speciations across phenotypic gradients; Fig 1B-D) have more severe effects on both model selection and parameter estimation (Figs. 4 and 5, TSS cases). For instance, increasing frequency of TSS in the simulations resulted in decreasing support for the BM model under a flat landscape, whereas models including the possibility of cladogenetic jumps were favored, namely the *κ* model and, to a lesser extent, the Lévy model. Under the TSS mode of speciation with normal landscape, phenotypic jumps at speciation also favored *κ* and Lévy models. In some cases, such jumps, appeared to disrupt the phylogenetic signal, with model selection providing support for the WN model, i.e. a model in which phylogenetic relationships do not contribute to explaining the trait values across species (Pennell et al., 2014; Silvestro et al., 2015). The lack of a predominantly favored model in TSS simulations with normal landscape reflects the lack of OU-like models with jumps. The increase support for WN models indicates a failure of macroevolutionary models to capture the true underlying process of evolution, which, in our simulations, is instead entirely linked to parent-offspring inheritance.

### Modeling cladogenetic jumps

While explicit models of cladogenetic and anagenetic evolution exist (Bokma, 2008), their implementation is currently unavailable. However, the *κ* transformation can capture cladogenetic change equally well as long as phylogenies are complete and in the absence of extinction (Harmon, 2018), which is the case in most of our simulations (Table A.1). The limitation of the *κ* model is that it does not explicitly model the jumps as a process and, as a consequence, does not estimate a corrected rate of anagenetic evolution. The Lévy process could fix part of the issue (Fig. 5B), but is unable to fully detect jumps at all nodes, because it is designed to pick up rarer events of evolutionary jumps (Duchen et al., 2017) instead of the more frequent events of cladogenetic jumps simulated here. Additionally, under the normal landscape, there are no implementations of models allowing us to jointly infer OU with jumps (an implementation of the model of Bartoszek, 2014, would prove ideal in this case). As a result, the best fitting model is often found to be a non-phylogenetic one (WN), indicating that current comparative methods fail to detect phylogenetic signal, which is blurred by jumps at speciation. Finally, both *κ* and Lévy do not exactly model the process that we simulated, but are preferred because of an artifact, which is the cladogenetic jumps generated by the natural sub-sampling of populations during a lineage split. In other words, where BM and OU models should be the best fit, TSS-induced jumps are blurring this signal. The same occurs under RSS if by chance the jumps are large enough to bias the process. There is thus a clear need for the development of new models that could account for such evolutionary processes.

### Punctuated equilibrium

Jumps at speciation events are predicted by “punctuated equilibrium”, a theory that postulates that most of the phenotypic change happens during cladogenesis (Gould and Eldredge, 1972, 1993). Since then, a major debate argued whether phenotypic change was predominantly anagenetic or cladogenetic (Uyeda et al., 2011), and new phylogenetic models to test these two hypotheses were, thus, developed (Pagel, 1999; Bokma, 2008; Futuyma, 2010; Bartoszek, 2014). This debate relies on adaptive jumps, while we were here concerned with jumps generated by sub-sampling of individuals at speciation events, which could be both adaptive or neutral. Consequently, while the Lévy jump model could detect evolutionary jumps at cladogenesis, it was not specifically designed to assess jumps at speciation events as it models the occurrence of a jump as an anagenetic process (Duchen et al., 2017; Landis and Schraiber, 2017). Additionally, the sub-sampling effect of populations on macroevolution is not restricted to speciation events only. Big changes in population sizes within branches of a phylogeny can also generate jumps or “bursts” in phenotypic traits (Uyeda et al., 2011), thus making this a more general issue.

### The next generation of evolutionary models for quantitative traits

While anagenetic evolutionary changes at the macroevolutionary level can be predicted by individual-based microevolutionary models (Rolland et al., 2018), it remains unclear how microevolutionary models would justify identical inheritance at speciation. These difficulties call for new models of macroevolution that account for cladogenetic change and flexibility concerning the directionality of phenotypic trajectories, i.e. flexibility concerning the fitness landscape of individuals in each species (similar to the model of Boucher et al. 2017, but allowing for asymmetric trait inheritance). Evolutionary models for discrete traits already account for non-identical inheritance at speciation, such as the “Dispersal-Extinction-Cladogenesis” (DEC) model (Ree et al., 2005; Ree and Smith, 2008), and the “Cladogenetic Stage change Speciation and Extinction” (ClaSSE) model (Goldberg and Igić, 2012). A continuous version of such models, that also include intra and interspecific variance (Kostikova et al., 2016), would allow us to infer the link between micro and macroevolutionary processes, which we formalized in the simulation framework presented here. Future directions might include studies that merge geographical and continuous trait models, which is now the case for biotic interactions among sympatric lineages (Nuismer and Harmon, 2015; Drury et al., 2016; Clarke et al., 2017; Quintero and Landis, 2019).

### Conclusion

We showed how different mechanisms of segregation of individuals into species at a microevolutionary time scale, affect overall macroevolutionary patterns across the entire phylogeny. The two modes of speciation we described here likely represent two extremes of a spectrum, and we expect that the true processes of inheritance of phenotypic traits will lie somewhere within that range. However, we argue that ecologically relevant or geographically differentiated traits will be generally subject to some level of segregation at speciation. Our results highlight the sensitivity of comparative methods to small variations in the inheritance of trait values between parent and daughter species, and call for caution when interpreting model testing and parameter estimation in macroevolution. In the light of the effects of differential trait segregation at speciation shown here, we propose the next generation of comparative methods incorporate asymmetric inheritance as a fundamental component of the evolutionary process.

## Acknowledgements

D.S. received funding from the Swedish Research Council (2015-04748) and from the Swedish Foundation for Strategic Research. N.S. received funding from the Swiss National Science Foundation (31003A-163428) and from the University of Lausanne. JR received funding from the European Union’s Horizon 2020 research and innovation programme under the Marie Skłodowska-Curie grant agreement No. 785910 and the Banting postdoctoral fellowship (151042). We thank the infrastructure of the Scientific Computing and Research Support Unit of the University of Lausanne (Switzerland) for the computational resources.

## Code and data availability

All the data and codes associated with this paper including meta-data and a tutorial to run the simulations are available at: https://bitbucket.org/daniele_silvestro/phylotraitsimulator/src/master/

## Supplementary Information

### A Supplementary tables

**Table A.1:** Number of extinct species per scenario. These statistics come from 100 simulated datasets per scenario.

| Scenario | Minimum | Median | Mean | Maximum |
| --- | --- | --- | --- | --- |
| RSS - flat | 0 | 0 | 0.42 | 3 |
| RSS - normal | 0 | 0 | 0.35 | 2 |
| Mixed 5% RSS - flat | 0 | 0 | 0.33 | 2 |
| Mixed 5% RSS - normal | 0 | 0 | 0.47 | 3 |
| Mixed 50% RSS - flat | 0 | 0 | 0.57 | 3 |
| Mixed 50% RSS - normal | 0 | 0 | 0.65 | 4 |
| TSS - flat | 0 | 1 | 0.81 | 6 |
| TSS - normal | 0 | 1 | 0.93 | 6 |

**Table A.2:** We assessed whether patterns of model testing under the different scenarios might be driven by differences in the relative branch lengths (all trees were rescaled to a depth of 1). We simulated traits under a BM model with rate parameter set to 0.01 along the trees generated by our simulations using the *fastBM* function (R package *phytools*, Revell (2012)). We then fitted BM, OU, *κ*, and White Noise (WN) models with *fitContinous* (R package *geiger*) and performed model testing using AICc scores. BM was predominantly selected as the best model across all simulations scenarios. These results show that the tree shapes and node depths of across scenarios do not influence model testing.

| Scenario | BM | OU | $\kappa$ | WN |
| --- | --- | --- | --- | --- |
| RSS - flat | 84 | 12 | 4 | 0 |
| RSS - normal | 83 | 10 | 7 | 0 |
| Mixed 5% RSS - flat | 86 | 10 | 4 | 0 |
| Mixed 5% RSS - normal | 84 | 11 | 5 | 0 |
| Mixed 50% RSS - flat | 80 | 16 | 4 | 0 |
| Mixed 50% RSS - normal | 87 | 9 | 4 | 0 |
| TSS - flat | 84 | 12 | 4 | 0 |
| TSS - normal | 83 | 9 | 8 | 0 |

### B Supplementary figures

**Figure B.1:**
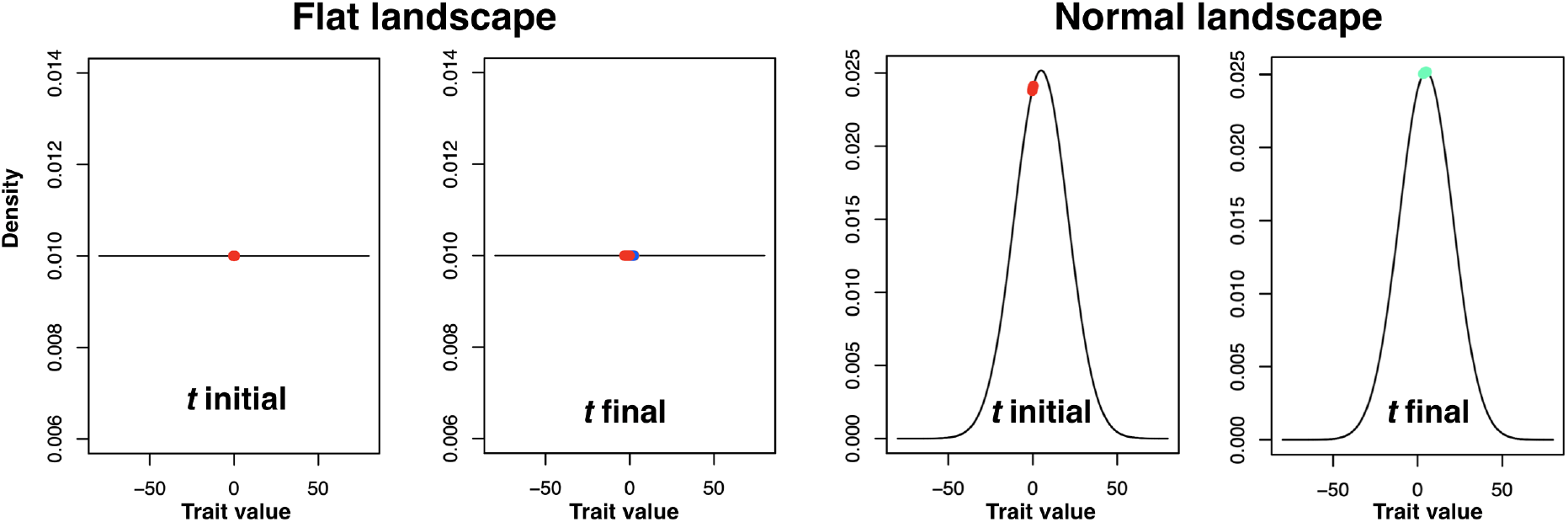
Fitness landscapes. Left: Flat landscape with the initial population (red points) and the set of species at the end of the simulation run (maintaining the same initial mean trait). Right: Normal landscape, where the final set of species has moved to the top of the landscape when compared to the initial values. 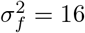 for the normal landscape of this particular example.

**Figure B.2:**
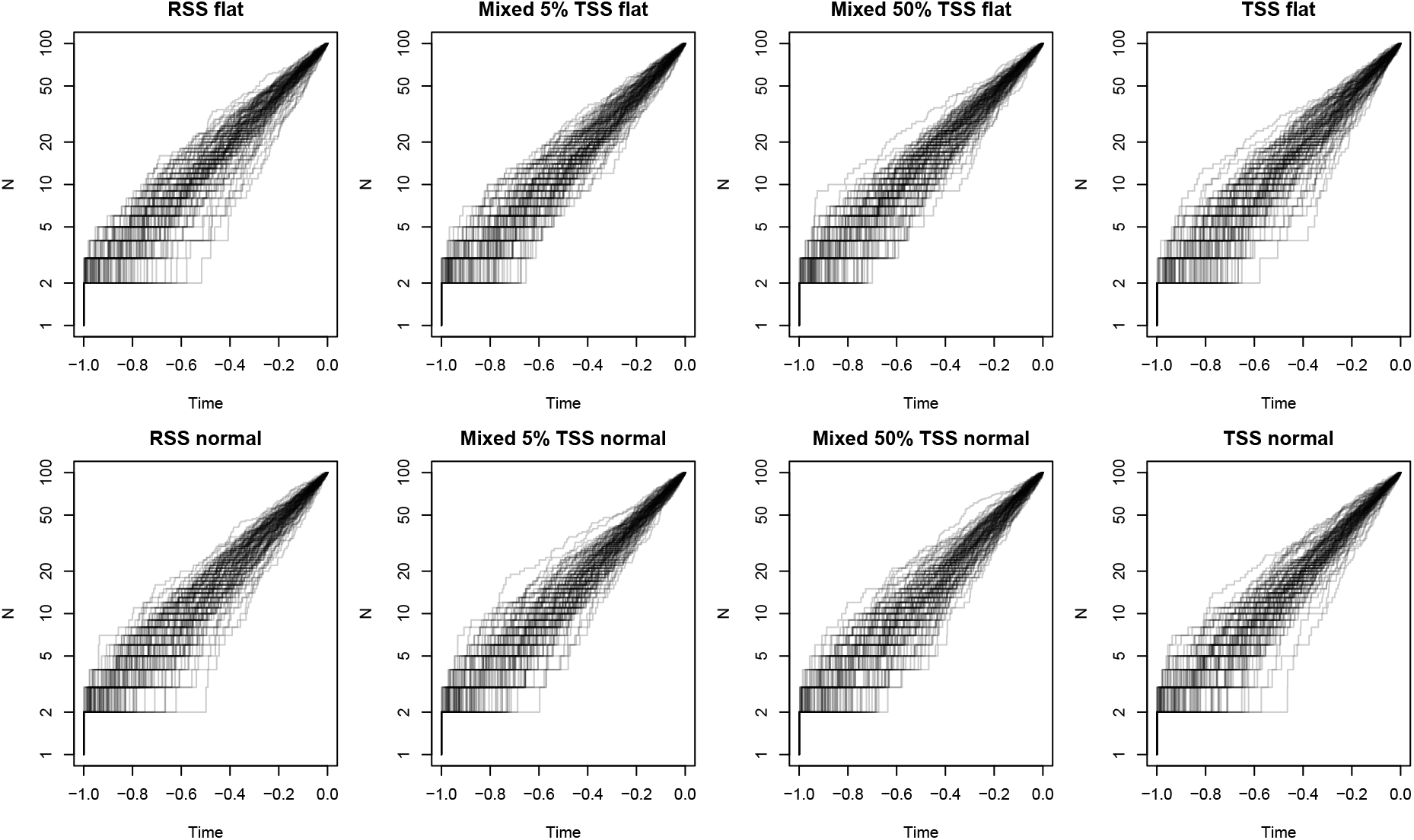
Lineage-through-time (LTT) plots of all simulations under all scenarios: RSS, mixed 5% TSS (95% RSS), mixed 50% TSS (50% RSS), and TSS for flat and (upper row) normal (lower row) fitness landscapes.

**Figure B.3:**
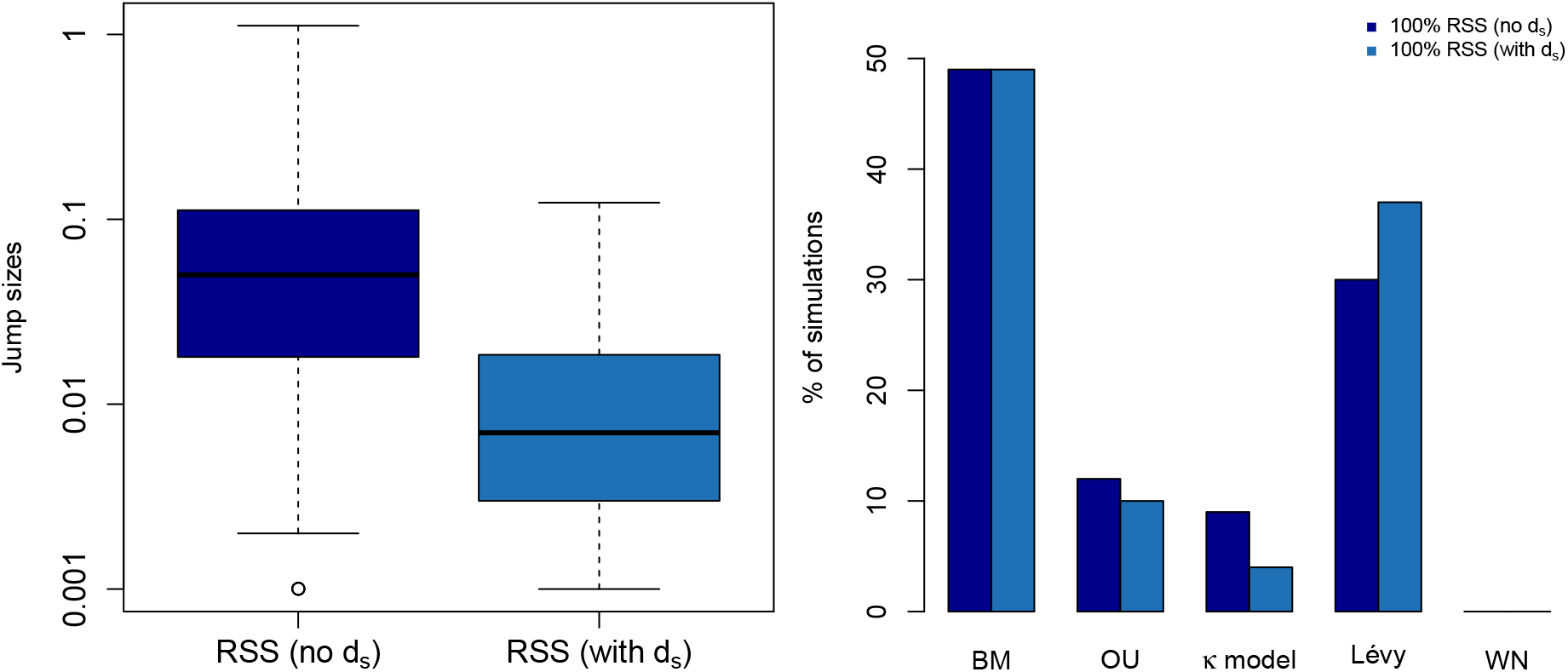
Absolute jump sizes (left) and model fit (right) for an RSS scenario where the death probability by distance to the species mean *d_s_* is not taken into account.

**Figure B.4:**
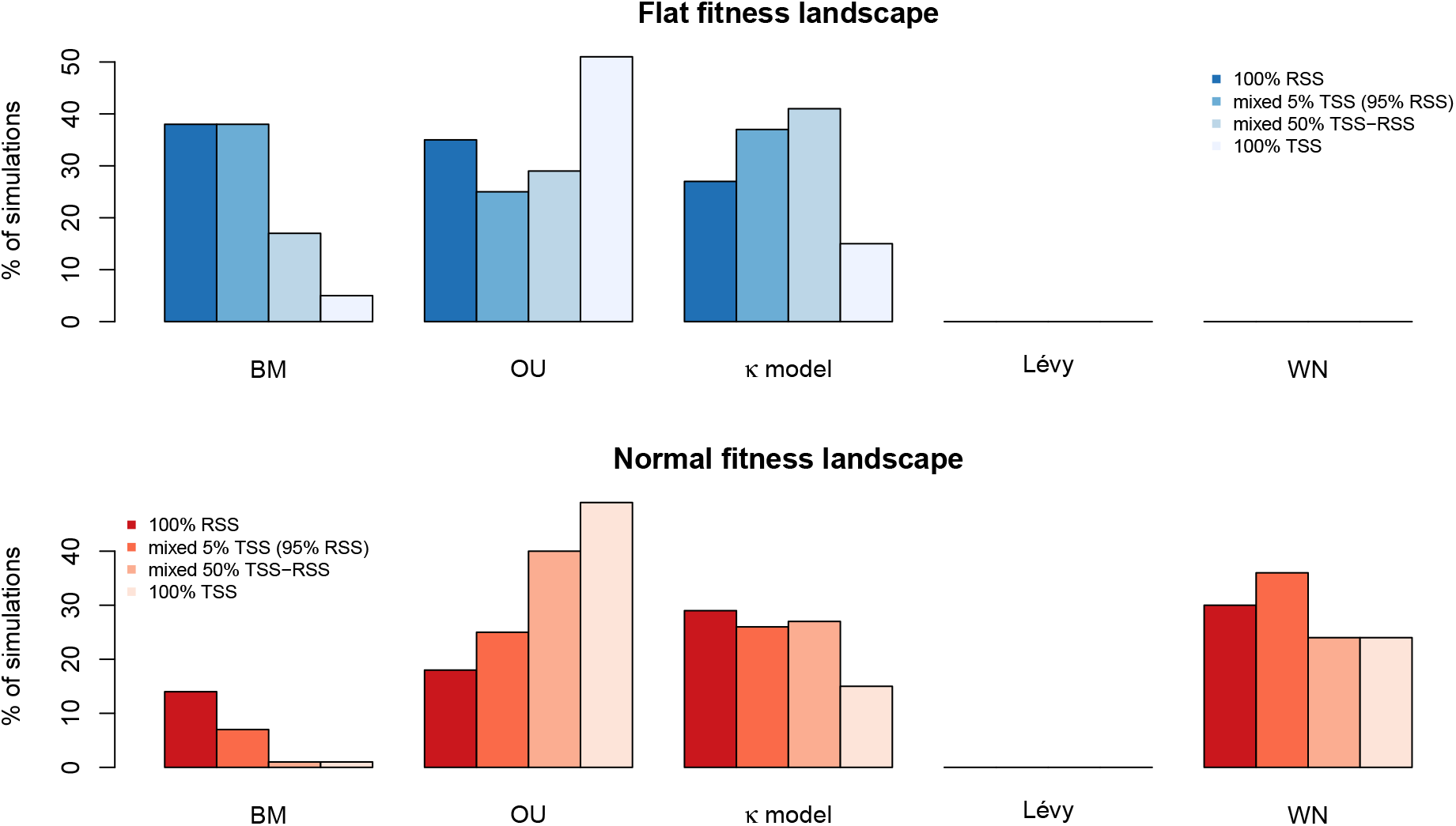
Second best fit under all scenarios. Bars represent the percentage of simulations that were second best fit by a given model. Abbreviations: BM = Brownian Motion, OU = Ornstein-Uhlenbeck, and WN = White Noise.

**Figure B.5:**
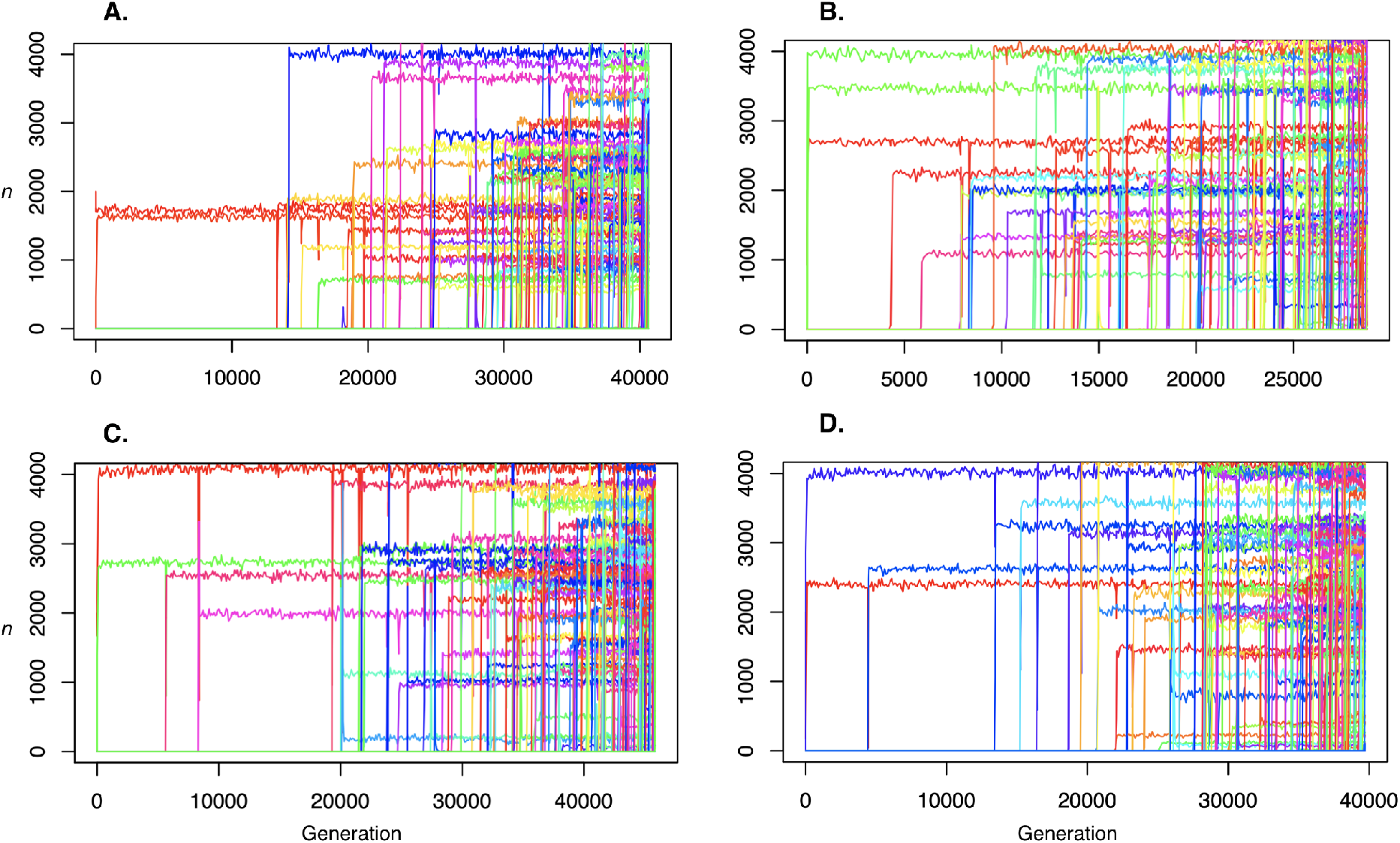
Change in population sizes *n* over time for the four extreme scenarios. A. RSS flat, B. TSS flat, C. RSS normal, D. TSS normal. Note how the carrying capacity *K* is reached very quickly.

## Notes

#### Summary of Updates

More simulation scenarios, including mixed RSS/TSS cases, and larger parameter ranges.

